# Stimulus-induced mechanical compaction of biological polymer networks via smart hydrogel microstructures

**DOI:** 10.1101/2025.04.01.646676

**Authors:** Vicente Salas-Quiroz, Katharina Esch, Katja Zieske

**Affiliations:** Molecular Biophysics and Living Matter, Max Planck Institute for the Science of Light, Erlangen, Germany; Department of Physics, Friedrich-Alexander Universität Erlangen Nürnberg, Erlangen, Germany

**Author notes:** Corresponding author (KZ), Max Planck Institute for the Science of Light, Staudtstr. 2, 91058 Erlangen, Germany.

**Keywords:** smart material, light-stimulus, thermoresponsive hydrogel, mask-less hydrogel patterning, NiPAAm, microstructures, extracellular matrix

## Abstract

The remodeling of the extracellular matrix by mechanical forces plays a crucial role in organizing cellular microenvironments. To study these mechanical perturbations, various methods have been developed to modify the cellular microenvironment and to apply controlled forces. However, most existing approaches rely either on instruments that cannot be integrated into lab-on-chip systems or on small probes with limited spatiotemporal precision. Here, we present a lab-on-chip system that enables spatially and temporally controlled mechanical perturbations of biological polymer networks. First, we fabricated thermoresponsive hydrogel microstructures within flow chambers and optimized their material composition and photopolymerization parameters. Second, we demonstrated the temporally controlled compression of Matrigel and collagen networks through temperature-stimulated expansion of the hydrogel microstructures. Following compression, Matrigel was plastically deformed, whereas the collagen network relaxed elastically. Finally, we showed that the compression of collagen networks can be spatially modulated by integrating hydrogel structures responsive to light stimuli. By mimicking the dynamic behavior of cells that remodel biological polymer networks, our method provides a versatile platform for future studies on extracellular matrix remodeling and the effects of mechanical forces on cellular microenvironments in both physiological and pathological contexts.

## Introduction

Mechanical remodeling of the extracellular microenvironment plays a crucial role in biological processes such as maintaining normal homeostasis (1), morphogenesis (2) and wound healing (3). Remodeling of the extracellular matrix is mediated by biochemical degradation or mechanical strains generated by cellular activity. Dysregulation of the extracellular matrix architecture or cellular activity is a hallmark of pathological conditions (4), such as cancer and fibrotic pathologies. For instance, tumor microenvironments exhibit increased mechanical activity exerted by cancer-associated fibroblasts (5). In addition, cancer cells have been shown to align collagen fibers and use this alignment to migrate more rapidly and persistently (6,7). Therefore, understanding how biological processes are impacted by mechanical remodeling of the extracellular matrix is essential for unraveling how the interactions of cells and their environment contribute to normal tissue development and pathological conditions.

Studying the influence of mechanical perturbations in living tissues systematically is challenging (8). Despite progress in optical methods such as multiphoton microscopy (9) and Brillouin measurements (10), living tissues are highly complex due to the large number of different cell types and their numerous interactions with extracellular components. To systematically investigate how mechanical perturbations affect biological processes, bottom-up approaches provide a promising strategy (11). Three-dimensional model tissues, such as organoids (12) and cancer spheroids (13) mimic specific properties of tissues or organs and provide more controlled biological systems. In addition, lab-on-chip systems are being developed to provide highly controlled fluid media compositions for culturing these model tissues and enabling parallelization (14).

However, how mechanical remodeling of microenvironments within lab-on-chip systems impacts the function and organization of three-dimensional model tissues remains poorly understood. One limitation of current techniques is the lack of suitable techniques to generate mechanical perturbation in biological polymer networks within lab-on-chip systems. Existing techniques for applying mechanical forces on cellular systems include atomic force microscopy, microneedles (15), and confinement force microscopy (16). However, these techniques are difficult to integrate into enclosed microfluidic lab-on-chip systems due to their large size and the need for direct contact with the samples. Alternative techniques, such as magnetic and optical tweezers (17), do not require direct contact with the sample and instead apply mechanical perturbation using small probes, which are guided by light or magnetic fields. However, these probes are typical spherical and the shape variations are limited. In addition, the number of independently controlled probes is often limited. Thus, the requirement arises for developing new microscale tools for mechanical perturbations of biological polymer networks that are compatible with lab-on-chip systems.

Smart hydrogels are powerful materials for microscale actuation and have been polymerized into diverse geometries (18). These hydrogels are composed of responsive polymers that change their structure in response to specific stimuli such as pH, light, or temperature (19). Thereby, smart hydrogel contracts or expands. We hypothesized that these actuations can be exploited to mechanically perturb biological polymer networks. While smart hydrogels have primarily been used in engineering sciences and studied for drug delivery and surface coating applications (20), their use in biomechanical applications remains limited. So far, light-sensitive hydrogel microactuators integrated with non-responsive components show promise for exerting forces on cells (21). In addition, smart hydrogel beads have been used to actuate cellular environments (22), and fillings of smart hydrogel have been used to actuate microfabricated pillars, which mechanically stimulated individual pillar-interacting cells (23). Moreover, light-responsive hydrogel sheets with grooves and coated with collagen have been employed to mechanically perturb cells plated on these surfaces (24). However, smart hydrogel microstructures with defined geometries have not yet been implemented for perturbation of biopolymer networks in lab-on-chip flow chambers.

To polymerize pre-gel solutions into defined geometrical microstructures various methods have been implemented. Standard photolithography enables the fabrication of hydrogel microstructures with defined two-dimensional geometries. However, this approach typically requires a photomask, limiting the geometries of hydrogel microstructures to the predefined features on the mask (25). Maskless methods for photopolymerization of hydrogel structures include two-photon lithography or light pattern modulation using a digital mirror device (DMD). Two-photon lithography allows for the fabrication of three-dimensional hydrogel shapes. However, this method is limited by its low speed and high cost (26). DMD-based devices, in contrast, define hydrogel microstructures in two-dimension. Compared to two-photon-lithography, their advantages include faster polymerization times and lower cost (27,28). Notably, many scientific questions related to perturbation, system development, and observation of minimal model tissues can be addressed in two-dimensionally defined systems (29). Thus, DMD-based polymerization of hydrogels represents a promising approach, integrating the flexibility of user-defined two-dimensional shapes with efficient fabrication times.

Here, we present an approach to remodel biological polymer networks with a high spatiotemporal control in flow chambers using smart hydrogels stimulated by temperature and light. We optimized polymerization parameters for generating hydrogel microstructures in flow chambers and demonstrated that the stimulated expansion of hydrogel microstructures leads to compaction of biological polymer networks, such as collagen and Matrigel. Moreover, we demonstrate that the mechanical remodeling of biological polymer networks can be highly controlled in space and time.

## Methods

### Reagents for thermoresponsive hydrogel

The thermoresponsive fraction of the pre-gel solution was composed 4.3 M Poly(N-Isopropylacrylamide) (NiPAAm) (Sigma Aldrich), 12 mM photoinitiator 2-Hydroxy-4’-(2-hydroxyethoxy)-2-methylpropiophenone (Irgacure 2959) (Sigma Aldrich), and 15 mM crosslinker N,N′-Methylenbisacrylamid (MBAm) (Thermo Scientific). NiPAAm and Irgacure 2959 were dissolved in ethanol. MBAm was dissolved in ultrapure water. The nonresponsive fraction of the pre-gel solution was composed of 4-arm poly (ethylene glycol)-acrylate (4-arm PEG) (MW10K, Laysan Bio Inc.) and the photoinitiator 4-Benzoybenzyl-trimethylammonium chloride (BOC Sciences). 4-arm PEG was dissolved in ultrapure water at stock concentration of 20%(w/v). 4-Benzoybenzyl-trimethylammonium chloride was dissolved in ethanol at a stock concentration of 100 mM. The stock solutions of 4-arm PEG, photoinitiator and water were mixed at a 1:2:1 ratio. The thermoresponsive and nonresponsive pre-gel solutions were mixed at various concentrations as described in the main text.

### Preparation of gold nanoparticles and light responsive hydrogel

The synthesis and coating of gold nanoparticles (AuNPs) followed the citrate reduction method (30). Briefly, gold(III) chloride hydrate (Sigma-Aldrich) was dissolved in ultra-pure water at a concentration of 1.2 mM in a piranha-cleaned Erlenmeyer flask. The solution was stirred and heated to 100°C. A solution of 170 mM sodium citrate tribasic dihydrate (Sigma-Aldrich) in ultra-pure water was added to the gold(III) chloride hydrate solution to a final concentration of 4 mM. The mixture was continuously stirred at 100°C for 30 minutes, whereby the color of the solution changed from pale yellow to dark red. The mixture was then cooled to room temperature.

To coat the AuNPs, 116 mM PEG_800_-SH (Sigma-Aldrich) in water was mixed with the AuNP solution at a 0.32:1 ratio and incubated at room temperature for 1 hour. The mixture was centrifuged at 21000 g for 15 minutes and washed with ultra-pure water. This washing step was repeated twice. Subsequently, the AuNP sediment (50 µl; original volume: 1ml) was collected and further concentrated by centrifugation. AuNPs in water were two-fold concentrated. To obtain AuNPs in ethanol, the nanoparticles were washed two times with ethanol and concentrated 16-fold.

For the preparation of the non-responsive pre-gel solution, a 20% 4-arm-PEG-acrylate solution was mixed with 4-Benzoybenzyl-trimethylammonium chloride (100mM), AuNPs (in water) and ultra-pure water at a ratio of 1:2:1. The light responsive pre-gel solution was composed of 4.3M NiPAAm dissolved in the AuNP (in ethanol) solution, 12 mM Irgacure 2959, and 15 mM MBAm. To obtain the final light-sensitive hydrogel mixture, the responsive and non-responsive pre-gel solutions containing AuNPs were combined at a 3:1 ratio. Pre-gel solutions with lower AuNP concentrations were prepared by mixing pre-gel solutions with and without AuNPs in varying ratios (Figure 4A-B).

### Flow chambers and temperature control

To fabricate flow chambers, two strips of Parafilm were used as spacers between two glass coverslips (22 mm × 40 mm and 22 mm × 22 mm). The assembly was heated on a hot plate at 200°C for 1 minute until the Parafilm melted. Pressure was applied with tweezers to achieve a channel height of approximately 0.1 mm.

For temperature control, a stage top incubator (Ibidi) was used. The incubator temperature was adjustable with an accuracy of ±2°C, and stability of the temperature was verified using a temperature sensor positioned in the center of the incubator. Alternatively, the flow chamber samples were heated by placing a glass-bottom petri dish (Ibidi) filled with water at 60°C (Figure 2) or 46°C (Figure 3G) on top of the flow chamber.

**Figure 1.**
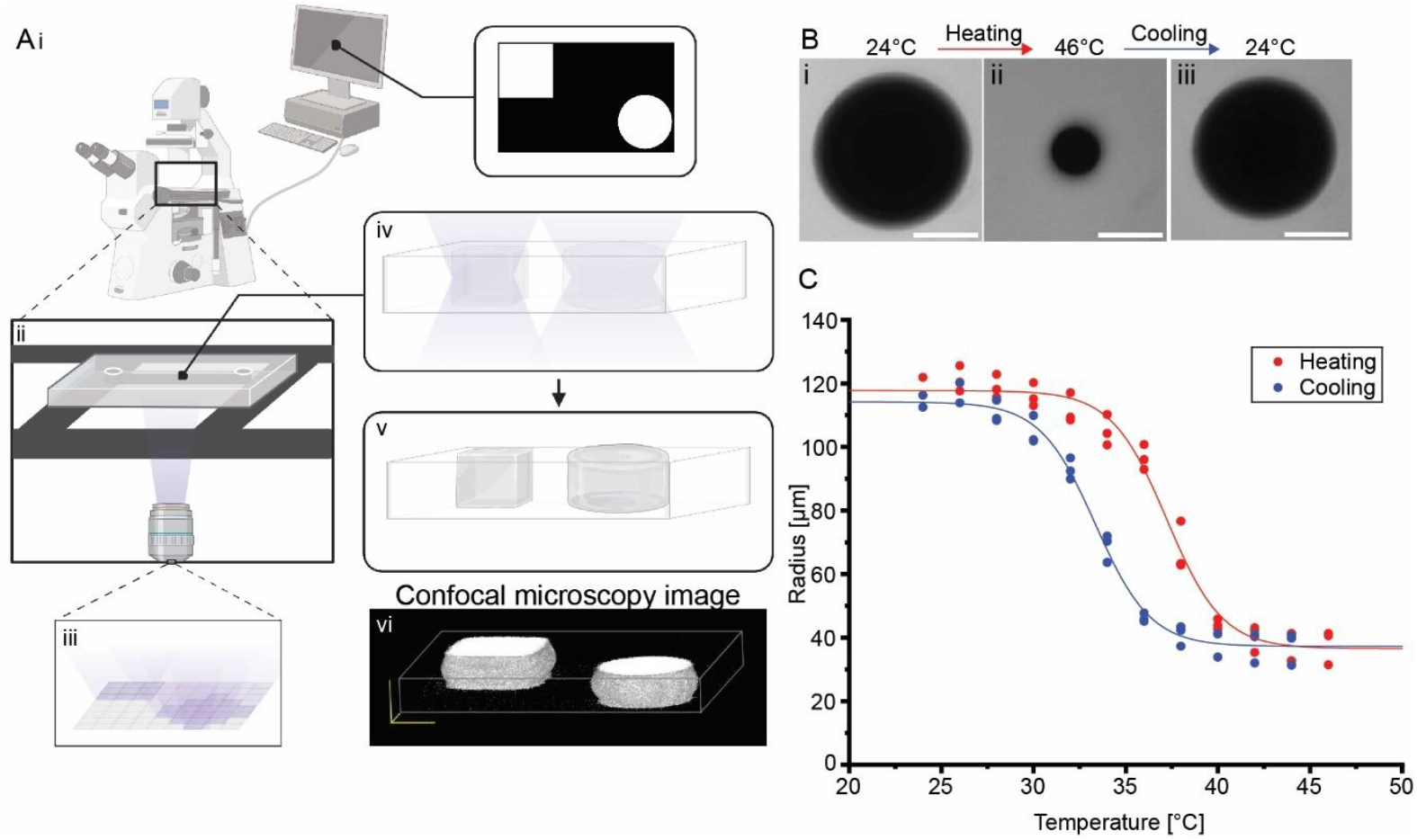
Workflow for mask-less photopolymerization of thermoresponsive hydrogel microstructures. **A**. Digital images with custom-designed patterns were interpreted by the software of a digital mirror device (PRIMO) connected to an inverted microscope (i). The hydrogel precursor solution within a flow chamber (ii) was exposed to the corresponding UV light patterns (iii, iv), resulting in the photopolymerization of hydrogel microstructures into defined shapes (v). (vi) A three-dimensional visualization of hydrogel microstructures reconstituted from z-stack images acquired using a laser-scanning confocal microscope. Created with Biorender.com. **B**. Confocal microscopy images of a cylindrical hydrogel microstructure at room temperature (24°C) (i), after heating to 46°C (ii), and after re-cooling to room temperature (iii). The flow chamber was supplemented with PLL-g-PEG/FITC (1 mg/ml). **C**. The cross-sectional area of hydrogel microstructures was quantified during heating (red) and cooling (blue) by measuring the radius. The sigmoidal curve was fitted using the dose-response equation. Heating curve: y = 36.6 + (117.8-36.6)/(1 + 10^((37.2-x)*-0.3)); Cooling curve: y = 37.2 + (114.2-37.2)/(1 + 10^((33.4-x)*-0.3)). Independent samples: n=3. Scale bars: 100 µm.

**Figure 2.**
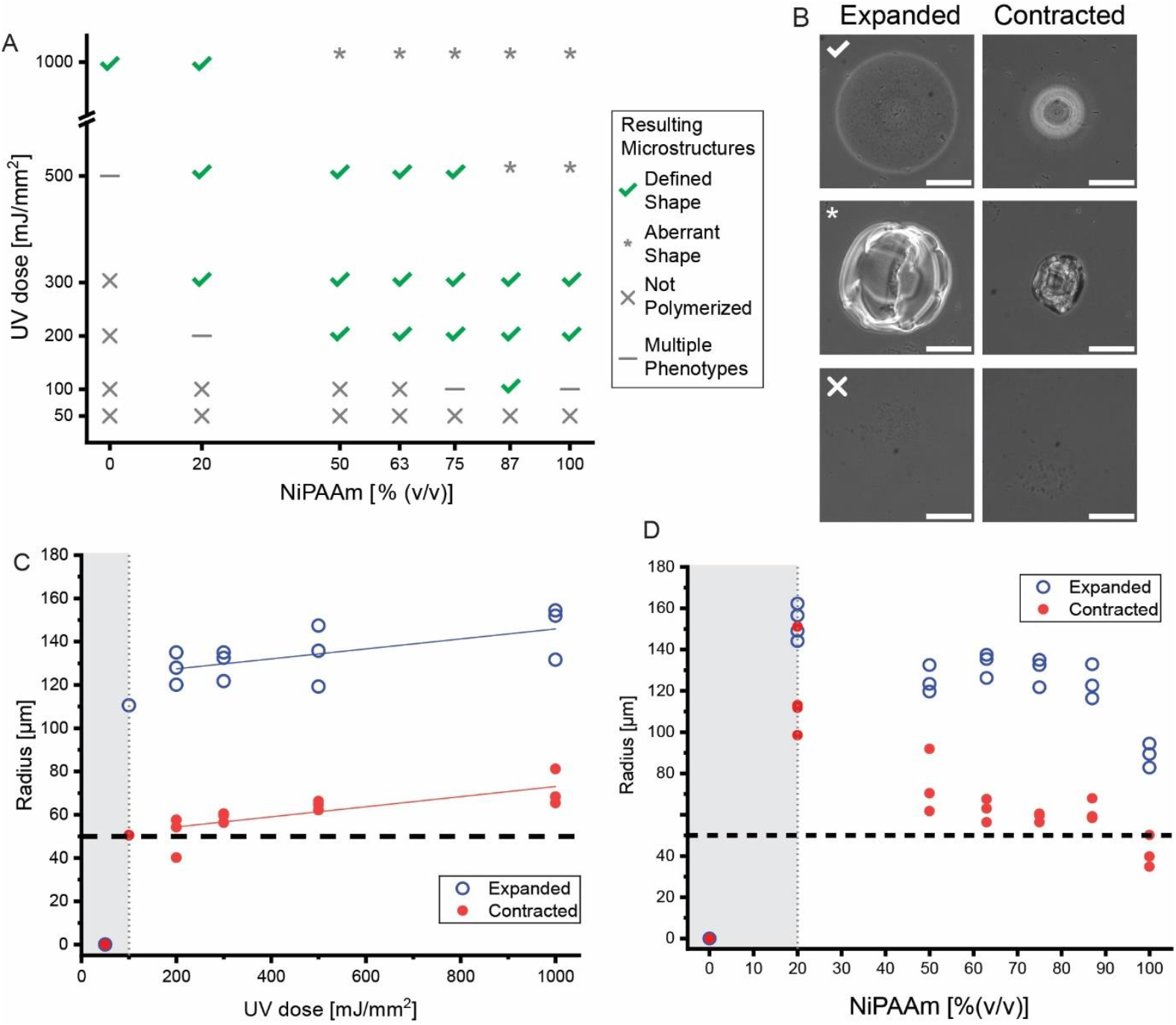
Optimization of illumination settings and material composition for generating thermoresponsive hydrogel microstructures. **A**. Hydrogel compositions with varying ratios of the NiPAAm-containing pre-gel solution (0% to 100%) were exposed to an UV light doses ranging from 50 mJ/mm^2^ to 1000 mJ/mm^2^. Depending on the specific combination of parameters, different phenotypes were observed: hydrogel microstructures did not polymerize (X), well-defined hydrogel microstructures that successfully polymerize (checkmark), multiple phenotypes appeared (-), and hydrogel microstructures with aberrant geometries polymerized (*). **B**. Representative images of the varioushydrogel microstructure phenotypes in their expanded state at room temperature and their contracted state at 60°C. Scale bars: 100 µm. **C**. Cross-sections of photopolymerized microstructures were measured as a function of UV dose. The fraction of NiPAAm-containing pre-gel solution was held constant at 75 % (v/v). The shaded grey area represents parameter combinations that did not result in the polymerization of defined structures. Data were fitted using the following functions: Expanded state: y = 122,84 + 0,02*x; contracted state: y = 49,69 + 0,02*x. **D**. Cross-sections of photopolymerized hydrogel microstructures were measured as a function of the fraction of NiPAAm-containing precursor solution. The UV dose was held constant at 300 mJ/mm^2^. The shaded grey area indicates parameter combinations that failed to polymerize well-defined structures. Independent samples for each condition (A-D): n=3.

**Figure 3.**
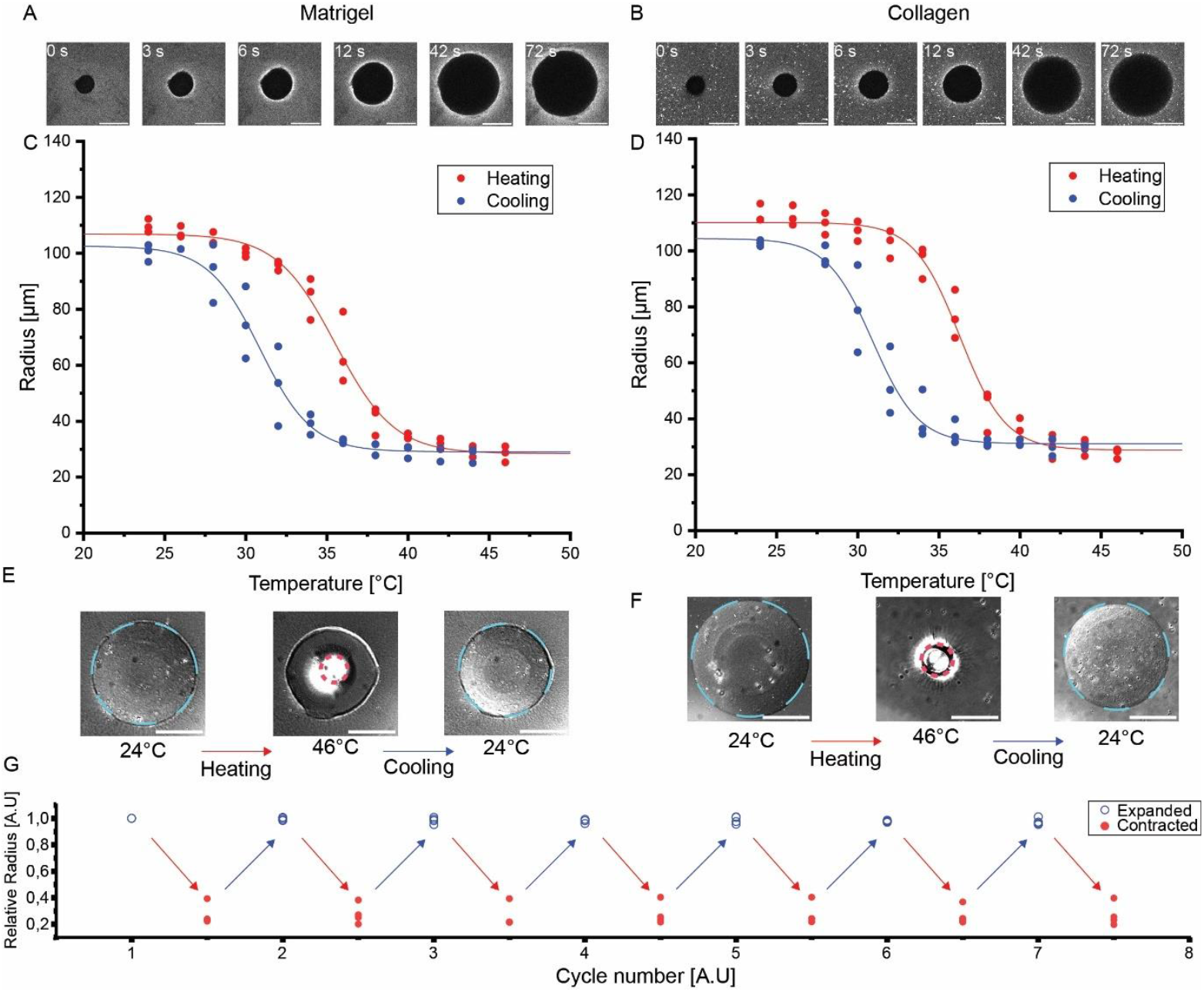
Biological polymer networks are mechanically compacted by expanding hydrogel microstructures. **A**. Confocal time-lapse microscopy images showing hydrogel microstructure expansion in the presence of Matrigel. To expand the hydrogel microstructure, the temperature was lowered from 37°C to room temperature (24°C). The Matrigel mixture was supplemented with the fluorescent probe PLL-g-PEG/FITC (47.6 µg/ml). **B**. The same experiment was performed in the presence of collagen. **C, D**. Quantification of the radii of cylindrical hydrogel microstructures during heating (red) and cooling (blue) in the presence of Matrigel (**C**) and collagen (**D**). Data were fitted with sigmoidal curves. Matrigel: heating curve: y = 28.8 + (110.1-28.8)/(1 + 10^((36.3-x)*-0.3)): cooling curve: y = 31.1 + (104.4-31.1)/(1 + 10^((30.9-x)*-0.3)). Collagen: heating curve: y = 28.4 + (106.9-28.4)/(1 + 10^((35.5-x)*-0.2)); cooling curve: y = 29.1 + (102.6-29.1)/(1 + 10^((30.9-x)*-0.3)). **E, F**. Representative brightfield images showing the state hydrogel microstructures within Matrigel and collagen at different temperatures during the heating-cooling cycle. The dashed circle indicates the boundaries of the microstructure. **G**. Quantification of multiple heating-cooling cycles demonstrating the repeatability of hydrogel structure size changes in collagen. Scale bars: 100 µm. Independent samples (A-F): n=3. (G): n≥3.

### Lithographic fabrication of hydrogel structures

Hydrogel microstructures were fabricated via maskless photolithography with a PRIMO device and Leonardo software (Alvéole, France). The PRIMO device was mounted on an inverted microscope (Nikon TI-E, Nikon Instruments) and projected patterns of ultraviolet light (wavelength: 375 nm) onto the flow chamber. Flow chambers were loaded with 15 µl of thermoresponsive or light-responsive hydrogel pre-gel mixtures and exposed to UV light patterns, while the central plane of the flow chamber was positioned in the focal plane. After exposure to UV light, the flow chambers were rinsed with 100 µl of ultrapure water and were used the same day.

### Sample preparation for fluorescent imaging and experiments with biopolymers

For imaging hydrogel structures in the presence of a fluorescent solution during temperature modulation experiments, the water was removed, and the flow chambers were filled with 15 µl of 1 mg/ml PLL-g-PEG/FITC (SuSoS). Both ends of the flow chamber were sealed with nail polish.

For experiments of hydrogel microstructures in Matrigel (Basement Membrane Matrix, Corning) or collagen (Type I, rat tail, Ibidi), water within the flow chamber was removed at 45°C and the chamber was cooled before filling with 15 µl Matrigel containing PLL-g-PEG/FITC (47.6 µg/ml) or a collagen solution. The collagen solution (pH=9.5, adjusted with sodium hydroxide) contained 4mg/ml collagen, 47.6 µg/ml PLL-g-PEG/FITC, 100 mM HEPES (VWR Chemicals), 3.7 g/l sodium bicarbonate, and ultrapure water. Both ends of the flow chamber were sealed with nail polish. Flow chambers with Matrigel were incubated at 37°C for 40 minutes. Those with collagen were incubated at 37°C for 15 minutes.

### Microscopy

Phase contrast images were acquired using a Nikon Ti-E microscope (Nikon Instruments) (Figure 2 and 3G).

Bright-field images were obtained using a Nikon Ti2 microscope (Nikon Instruments) equipped with a 20x objective (CFI Plan Fluor 20x, Nikon Instruments) (Figure 4A, B). A 460nm LED (power: 259 mW) was used to trigger volume change of the light-responsive hydrogel structures.

Confocal images and z-stacks were acquired using a confocal laser scanning microscope (LSM 980, Zeiss) equipped with an 10x objective (Plan-Apochromat 10x/0.45 M27, Zeiss) (Figure 1 and 3A-F). For spatially selective hydrogel microstructures contraction (Figure 4 C-E), an area of interest was illuminated with a 533 nm laser.

**Figure 4.**
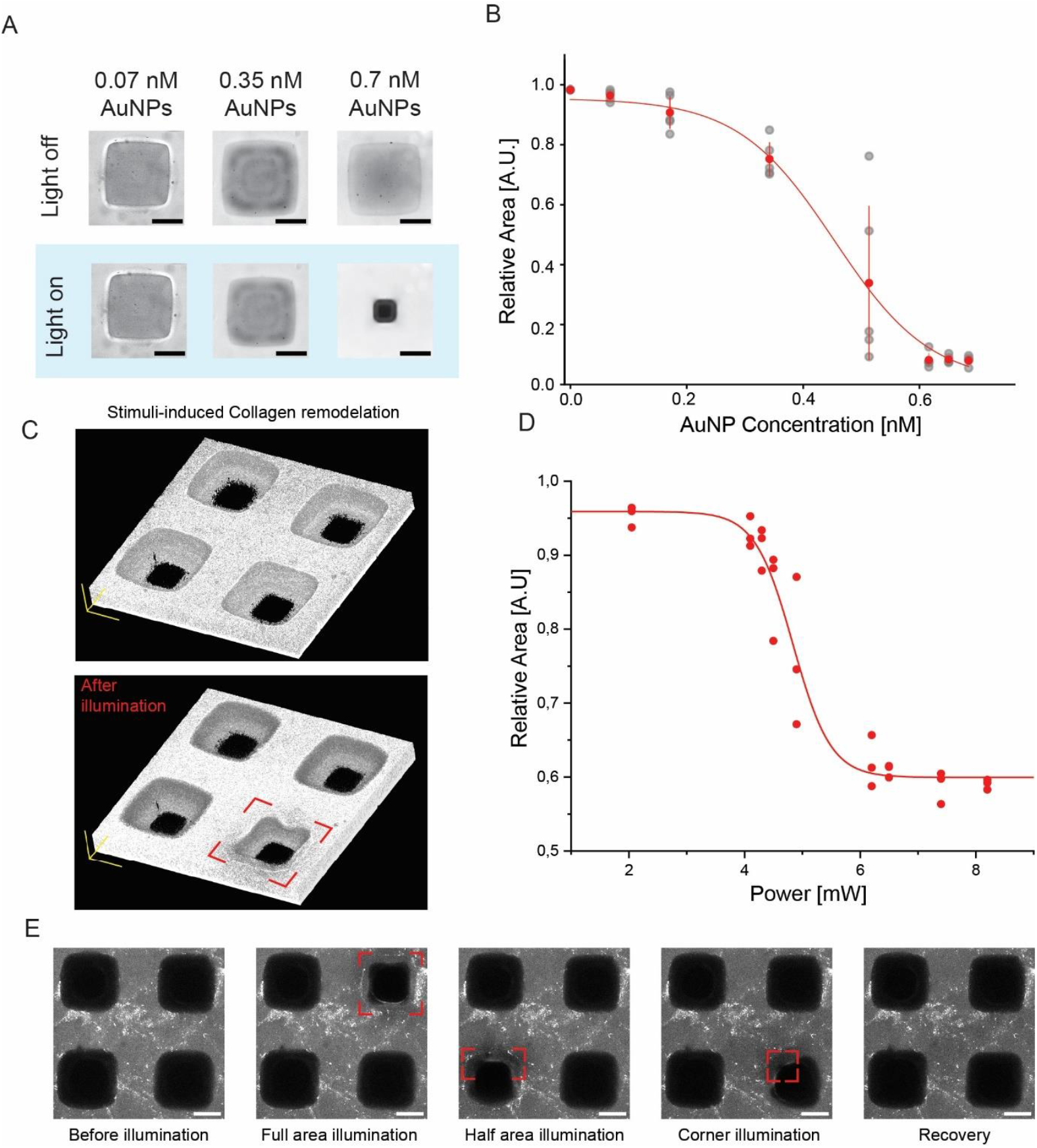
Collagen deformation through light-stimulated contraction of individual hydrogel blocks. **A**. Representative images of hydrogel microstructures with varying AuNP concentrations before and after light exposure (460 nm, 259 mW) for 3 minutes. **B**. Relative area change of the hydrogel microstructures as a function of AuNP concentration. Error bars indicate the standard deviation. Fit function: y = 0.97/(1+e^(11.81*(x-0.46)) **C**. Three-dimensional reconstruction of collagen sample before and after illumination, aquired using a laser-scanning confocal microscope. The red square marks the Illuminated hydrogel microstructure. **D**. Quantification of light-driven compaction of hydrogel structures as a function of laser power. Fit function: y = 0,6 + (1-0,6)/(1 + 10^((4,8-x)*-1,3)). **E**. Representative images of light-responsive hydrogel microstructures showing a microstructure before illumination, compaction upon full or subregional illumination (indicated by red squares), and recovery after cessation of light exposure. Independent samples (A-B): n ≥3, (C-E): n=3. Scale bars: 100 μm.

### Software for Data Analysis

Digital masks (TIFF images, 8-bit, 0.27 µm/px) were designed using the software Inkscape.

Hydrogel geometries in phase contrast images were analyzed using Fiji (31). The analysis to retrieve the area of the imaged microstructures was performed on manually drawn masks along the boundaries of the hydrogel microstructures.

For confocal images, Fiji was used to retrieve the radius of the microstructures. Masks were generated via thresholding on the central z-stack slice.

Origin (OriginLab) and Python were used for plotting and statistical analysis.

## Results and Discussion

To exert forces on biological polymer networks, we established a workflow for fabricating thermoresponsive hydrogel microstructures via mask-less photoprinting and optimized the printing parameters. The pre-gel solution was composed of two photoinitiators - Irgacure 2959, which enables free radical polymerization (32), and 4-Benzoybenzyl-trimethylammonium chloride – along with the crosslinker MBAm, the non-responsive 4-arm-PEG and the thermoresponsive polymer NiPAAm. Previously, we demonstrated that a pre-gel solution containing these components adheres to glass surfaces and expanded upon temperature stimuli (33). The liquid pre-gel mixture was injected into a custom-made flow chamber. To photopolymerize the precursor solution into defined microstructures, patterns of UV light were projected onto the sample using a digital mirror device (PRIMO, Alvéole) (Figure 2A). By generating circular and rectangular shapes, we confirmed that different custom-defined microstructure geometries could be fabricated. (Figure 1Ai-vi). Compared to our previous study, where photomasks were applied (33), the current setup allows for the fabrication of hydrogel microstructures with any custom-defined 2D geometry. In addition, the corresponding software was calibrated to adjust illumination times and achieve controlled illumination doses.

After polymerization of the hydrogel microstructures, the thermoresponsiveness was confirmed by heating and cooling the hydrogel microstructures within an incubator chamber. To visualize hydrogel boundaries, a fluorescent probe was used that did not penetrate the hydrogel blocks. Laser scanning confocal imaging was used to acquire z-stacks and visualize the boundaries of the hydrogel structures (Figure 1Avi), that were easily distinguishable from their environment (Figure 1B). When the temperature was increased to 46°C, cylindrical hydrogel microstructures shrank to approximately 20% of their original size at room temperature (24°C). This contraction was reversible upon cooling (Figure 1B).

The volume phase transition temperature (VPTT) for NiPAAm hydrogels has been determined to be around 33°C (34,35). To quantitatively analyze the VPTT of our hydrogel microstructures and determine the size changes as a function of temperature, we varied the temperature in increments of two degrees between 24 and 46°C (Figure 1C). The data for cooling and heating cycles were fitted with a sigmoidal curve, respectively. Our data verified the presence of thermal hysteresis, which has been reported previously (36,37). The VPTT of our hydrogel microstructures during the heating cycle was 37.2±2°C. During the cooling cycle, the VPTT was 33.3±2°C. These VPTTs have advantages for future cellular applications, as size changes can be triggered within the physiological relevant temperature range (38,39). The expansion of these fabricated hydrogel microstructures therefore provides an intriguing opportunity for applying pushing forces in biological contexts within lab-on-chip chambers.

Next, we determined the optimal illumination dose and material composition for microstructure fabrication and controlled gel expansion. Previously, we reported a hydrogel mixture for generating adherend thermoresponsive microstructures on glass (33), which consisted of a thermoresponsive pre-gel fraction including NiPAAm and a non-responsive material fraction, containing 4-arm PEG. Here, we systematically studied how varying the ratio of the NiPAAm containing fraction (0, 20, 50, 62, 75, 87, and 100% v/v) and adjusting the exposure time affected the formation of hydrogel microstructures (Figure 2A). Thereby, we observed three major outcomes. First, at low UV doses and low fractions of the NiPAAm containing pre-gel solution, the hydrogel did not polymerize. Similarly, at a very low UV dose of 50 mJ/mm^2^, even high NiPAAm-containing formulations did not polymerize, indicating that this energy level was insufficient to initiate polymerization (Figure 2A, 2B bottom). Second, at medium UV doses and medium-to-high fractions of the NiPAAm containing pre-gel solution, the hydrogel polymerized into geometrically well-defined and thermoresponsive microstructures (Figure 2A, 2B, top). Finally, at high UV doses and high fractions of the NiPAAm containing pre-gel solution, the resulting microstructures exhibited aberrant geometries (Figure 2A, 2B, middle).

Next, we quantified the thermoresponsiveness of the geometrically well-defined hydrogel microstructures by measuring their radius in the expanded and contracted states. First, we measured the dependence of the microstructure radii on UV dose, using a fixed fraction of the NiPAAm containing pre-gel solution of 75% (w/v) (Figure 3C). For UV doses between 200 mJ/mm^2^ and 500 mJ/mm^2^, the hydrogel microstructures exhibited well-defined geometries and displayed clear temperature responsiveness. The radius of the hydrogel microstructures increased with increasing UV doses. This trend is likely caused by prolonged light exposure resulting in polymerization beyond the intended pattern areas.

We also measured how the fraction of the NiPAAm containing pre-gel solution affected the microstructure radii while maintaining a constant UV dose of 300 mJ/mm^2^ (Figure 3D). Without NiPAAm, no hydrogel microstructures polymerized. At low NiPAAm fractions, the resulting structures were larger than the digital templates. Geometrically defined hydrogel microstructures, which displayed clear temperature responsiveness, were observed for NiPAAm containing fractions above 50%.

Together these results establish a mask-less approach for patterning thermoresponsive hydrogel microstructures within flow chambers. We found optimal fabrication parameters that ensure microstructures reflect the geometry of the custom-designed digital templates while maintaining responsiveness and adhesion to the flow chamber surface. The optimal fabrication parameters were identified to be a UV dose of 200 mJ/mm^2^ - 500 mJ/mm^2^ and a NiPAAm containing pre-gel solution of 50-100% v/v. For subsequent experiments, we used a UV dose of 300 mJ/mm^2^ and a NiPAAm containing pre-gel solution of 75% v/v. Under these conditions, the hydrogel microstructures exhibited controlled size changes, enabling precise application of mechanical forces to their surroundings.

Biological tissues are dynamic, frequently undergoing movement and remodeling due to the interaction between cells and their environment (40). Cellular environments are often composed of biological polymers, that may elastically or plastically respond to mechanical perturbation and forces (41,42). Therefore, developing systematic approaches for mechanically remodeling biological polymers networks would be promising for studying the underlying mechanisms of tissue dynamics. We investigated whether expanding hydrogel microstructures could serve as microscale tools for studying the effect mechanical perturbations on biological polymer networks. By embedding our hydrogel microstructures in extracellular matrix polymer networks, we aimed to expose these polymer networks to mechanical strains and forces. Specifically, we filled the hydrogel structure containing flow chambers with Matrigel or collagen. After gelling, the samples were subjected to heating and cooling cycles and imaged to study the behavior of the microstructures and the surrounding polymer networks.

First, we characterized samples containing membrane-based Matrigel, a widely utilized extracellular matrix-based material for mimicking cellular environments. For visualization purposes, we supplemented the Matrigel with FITC-labeled PLL-g-PEG. The Matrigel was gelled within the flow chamber at 37°C while the hydrogel microstructures were in the contracted state. Upon cooling to room temperature, the hydrogel microstructures expanded and pushed against the surrounding Matrigel. This expansion caused a compaction of the Matrigel (Figure 3A). This demonstrated that our microstructures generate sufficient force to mechanically remodel the gelled Matrigel network. The local compaction was further evidenced by the accumulation of the fluorescent probe around the rim of the microstructure during expansion (Figure 3A). Compared to a sample without Matrigel, the expanded hydrogel microstructures in Matrigel were smaller. This size difference in aqueous solution and within Matrigel is likely caused by the mechanical resistance of the surrounding Matrigel network. After the initial expansion, the samples were heated to 46°C and subsequently cooled to 24°C (Figure 3C, E) to characterize the volume dynamics of the hydrogel structures. The sigmoidal compaction and expansion dynamics of the microstructures were similar in the presence and absence of Matrigel.

Notably, during hydrogel contraction, a gap appeared between the Matrigel and the microstructure (Figure 3E, middle panel). This suggests that Matrigel deforms plastically under mechanical perturbations imposed by expanding hydrogel structures. This observation aligns with previous studies reporting plastic remodeling of Matrigel (43–46) and demonstrates the potential of smart hydrogel microstructures as tools for generating localized, plastic deformations in Matrigel at the micron scale. In vivo, the plastic remodeling of extracellular polymer networks by cellular forces plays a crucial role in processes such as cancer cell invasion of basement membranes (42).

To characterize another biological polymer network within our system, we filled the flow chambers with type I collagen, a widely studied biopolymer in biomedical and bioengineering research (47). Similar to Matrigel, we observed compaction of the collagen network during the initial expansion of the hydrogel microstructures (Figure 3B). However, in contrast to Matrigel, the collagen network behaved elastically upon contraction of the hydrogel microstructures (Figure 3F, middle panel). The collagen network dynamically refilled the space vacated by the contracting microstructures, demonstrating that our system cannot only be applied to induce pushing forces, but also to allow the relaxation of collagen networks.

We performed multiple heating-cooling cycles with collagen samples (Figure 3G) and demonstrated that the compaction and expansion of collagen networks were reversible and reproducible. After seven heating and cooling cycles, the radii of the microstructures in both the expanded and compacted states, did not vary more than 5%. This demonstrates the long-term stability and usability of the thermoresponsive hydrogel structures as micron-scale tools for generating mechanical forces in biopolymer networks. Our assay represents a novel biomechanical approach for remodeling biopolymer networks in lab-on-chip systems. Due to the fibrillar nature of collagen (48), which facilitates force transmission, deformations induced by the hydrogel structures could be detected several micrometers away from the source.

After demonstrating network compaction around microstructures in response to an external temperature stimulus, we further functionalized our hydrogel assay to be responsive to light stimuli. We aimed to use light as a trigger for inducing size changes of our hydrogel microstructures for three reasons: First, light can be adjusted to a target intensity more rapidly than an incubation chamber can reach a target temperature, allowing for faster switching. Second, while an entire sample is exposed to heating within an incubation chamber, a laser beam can be spatially guided, enabling localized control of hydrogel compaction and expansion. Finally, many biological samples are temperature sensitive. Using spatially controlled light patterns to stimulate localized temperature changes only within the hydrogel microstructures, while minimizing heating of the surrounding biological polymer network, could expand applications to temperature-sensitive samples.

To render hydrogel microstructures light-responsive, gold nanoparticles (AuNP) have previously been integrated into hydrogels to enable plasmonic heating upon light exposure (23,30). To determine a AuNP concentration sufficient for inducing light-dependent volume changes in our hydrogel microstructures, we systematically varied the AuNP concentration in our pre-gel solution. Pre-gel solutions containing distinct concentrations of AuNPs were patterned into square microstructures and afterwards stimulated with an LED (460 nm, 259 mW) for 3 minutes. Imaging the hydrogel microstructures before and after stimulation demonstrated that light-induced compaction was more pronounced at higher AuNP concentrations compared to low amounts of AuNP (Figure 4A). We quantified the relative areas of the expanded and compacted state of the hydrogel microstructures for different AuNP concentrations. The data were fitted with a sigmoidal curve that exhibited an inflection point at 0.45 nM AuNPs (Figure 4B). This result demonstrated that AuNP concentrations above 0.45 nM effectively modulate hydrogel size changes. For the following experiments, we used a concentration of 0.63 nM AuNP.

Next, we tested whether light patterns could stimulate the contraction of individual hydrogel microstructures embedded in a collagen matrix. To do this, arrays of AuNP-containing hydrogel microstructures were generated and embedded in 4 mg/mL collagen. We selectively exposed a single hydrogel microstructure to light (561 nm) using a laser scanning microscope and observed contraction of the targeted hydrogel microstructure (Figure 4C). The laser intensity was systematically varied and the compaction of the hydrogel was quantified. The data revealed a sigmoidal response, similar to temperature-dependent contraction, with higher laser intensities resulting in more pronounced compaction (Figure 4D). These results demonstrate the controlled expansion of individual hydrogel microstructures within our flow chambers.

We further explored spatial control of our samples by selectively illuminating subregions of the microstructures (Figure 4E). For instance, we exposed half of a square hydrogel microstructure and then a quarter of microstructure to light. We observed contraction in the illuminated regions, while the non-exposed regions did not contract. These deformations were reversible, with hydrogel microstructures recovering their original shape and size upon cessation of illumination. The ability to control the contraction of specific subareas of the microstructures in both space and time illustrate the high level of control in hydrogel shape modulation and the resulting mechanical remodeling biological polymer networks.

In summary, this work presents a novel approach for remodeling biological polymer networks using smart hydrogel microstructures. The expansion and compaction of these hydrogel microstructures were triggered by either light or temperature providing high spatiotemporal control over mechanical perturbations of biological polymer networks, such as collagen and Matrigel. Our approach is well-suited for lab-on-chip applications, with potential future applications including the characterization of diverse extracellular matrix compositions and the mechanical perturbation of cellular systems.

## Acknowledgements

We thank all members of the Zieske research group for scientific discussions. Language and grammar of the manuscript was refined with assistance of Chat GPT (v3.5)

